# Brain digital twins reveal network changes in congenital and slowly progressive cerebellar ataxias

**DOI:** 10.64898/2026.03.23.713380

**Authors:** Marta Gaviraghi, Anita Monteverdi, Sara Bulgheroni, Marta Mercati, Arianna De Laurentiis, Anna Nigri, Marina Grisoli, Stefano D’Arrigo, Claudia AM Gandini Wheeler-Kingshott, Claudia Casellato, Fulvia Palesi, Egidio D’Angelo

## Abstract

Cerebellar ataxias are a rare group of disorders manifesting with motor incoordination and cognitive-affective deficits of variable severity. Although neurogenetic has revealed multiple mutations, the study of ataxias still relies on clinical evaluation, while the underlying neural network changes remain unclear. It has been argued that the less severe symptoms in congenital (like Joubert syndrome, JS) than in slowly progressive (SP) ataxias reflect a different interplay of alteration and compensation but direct evidence is still lacking. Moreover, it is unclear why, in front of a wide heterogeneity of molecular alterations, SPs show common clinical symptoms.

To address these questions, we created brain digital twins for each participant by combining volumetry, graph theory analysis of structural and functional connectivity, and dynamical simulations using the virtual brain. We studied 8 JS (3 females, 21±6years), 8 SP (3 females, 20±5years) and 11 healthy controls (HC; 5 females, 21±2years).Volumetry quantified atrophy, graph metrics (centrality, segregation and integration) characterized topology, and neural dynamical simulations estimated excitation/inhibition balance, providing anatomo-physiological parameters within the somatomotor (SMN) and ventral attention (VAN) networks. Anatomo-physiological parameters were correlated with clinical/neuropsychological scores, and unsupervised clustering was applied to assess whether network features can discriminate between JS and SP beyond clinical classification.

MRI morphometry confirmed selective vermis reduction in JS and a widespread cerebellar atrophy in SP compared to HC. In both ataxia groups, SMN and VAN showed reduced volume and structural connectivity but with different patterns of topological and dynamical alterations. In the SMN of SP, reduced centrality and excitation/inhibition balance depressed information transfer through the network. In the VAN of JS, reduced centrality, segregation, and integration, were detrimental but coexisted with a higher number of functional core nodes and an increased large-scale excitatory coupling, supporting compensatory reorganisation in extracerebellar nodes. Clustering confirmed that SMN better differentiates SP, whereas VAN better clusters JS. Importantly, anatomo-physiological parameters of network volume, topology, and dynamics correlated with patients’ motor and cognitive performance.

In conclusion, primary cerebellar damage secondarily impacts large-scale brain networks, altered in both ataxia groups but compensated only in JS. Similar clinical symptoms in SP reflects the similarity of network changes, while differential involvement of SMN and VAN in JS and SP reflects the connectivity pattern of the lesioned areas inside these large-scale brain circuits. Importantly, anatomo-physiological parameters are sufficient to explain individual motor and cognitive performance, offering a basis for improved patient profiling and personalized therapies.

## Introduction

Cerebellar ataxias are a group of disorders primarily affecting the cerebellum, characterized by impaired balance and coordination in the absence of muscle weakness^1^. Cerebellar ataxias can be categorized into non-progressive congenital forms that remain stable over time, such as Joubert syndrome (JS), and hereditary or metabolic slowly progressive (SP) cerebellar ataxias, which are marked by gradual cerebellar atrophy and progressive worsening over years or decades^2^. Congenital ataxias are clinically characterized by variable degree of motor developmental delay, very early-onset cerebellar ataxia, and variable cognitive impairment. Patients typically follow a non-progressive course, with most showing gradual improvement in motor and cognitive skills over time^3,4^. Conversely, SP cerebellar ataxias involve a progressive decline in motor function, with cognitive deterioration occurring in certain forms, possibly due to the impossibility to generate effective compensation under the continuous action of the etiopathogenetic causes^4,5^.

JS is a rare genetic neurodevelopmental disorder, with an estimated prevalence from 1 in 80000 to 1 in 100000 live births^6^. The hallmark, visible on MRI scans, is the “molar tooth sign”, already evident at birth and characterized by a deep interpeduncular fossa, enlarged superior cerebellar peduncles, and hypoplasia of the cerebellar vermis^7,8,9^. Despite marked genetic heterogeneity^10,11,12^, JS shows a rather stereotyped pattern of alterations of cerebellar circuit formation and morphogenesis^13,14,15,16,17^. In contrast, SP ataxias encompass a wide spectrum of genetic and metabolic disorders characterized by generalized cerebellar atrophy progressing gradually with childhood onset. These include spinocerebellar ataxias^18,19,20^, metabolic deficiencies^21^, and complex multisystem disorders^22^. Despite their diverse causes, SP share common features such as cerebellar maldevelopment, progressive cerebellar atrophy, impaired motor coordination and cognitive-affective functions. Beside neurogenetics and anatomo-clinical correlations, several questions remain open. What differentiates SP from JS in terms of network properties? What are the presumed network compensation mechanisms taking place in the JS brain and reducing the gravity of symptoms? Is there a common pattern of network alterations in the SP brain despite the diversity of etiopathogenetic causes? Answering these questions implies considering the cerebellar regions as part of specific functional networks, rather than focusing on local cellular and anatomical alterations^23^.

Despite the remarkable progress on the molecular, cellular, and microcircuit physiology of the cerebellum^24,25^, very little is known about network alterations leading to cerebellar ataxia. To date, literature focused mainly on qualitative cerebellar morphometry^26,27^ and microstructural analysis^28^. Here, we combined measurements of regional volumes, graph theory analysis and computational simulations of brain function using virtual brain models (TVB) to uncover the relationship between structure, function, and dynamics^29^ in the ataxic brain. This approach enabled the construction of subject-specific brain digital twins, capturing network specific anatomo-physiological parameters. Graph theory enables the characterization of topological properties of structural and functional networks identified by diffusion and functional MRI data^28^. TVB allows subject-specific simulations of brain activity by integrating structural connectome data with region-specific dynamic equations (e.g., the Wong-Wang model), which describe large scale signals in terms of the interaction between excitatory and inhibitory populations^30,31,32^. We focused on two networks including the cerebellum as a key node: the somatomotor network (SMN), since motor dysfunction is the main clinical manifestation in subjects with cerebellar ataxias^33,34,35^; and the ventral attention network (VAN), because of the increasingly recognized role of the cerebellum in attention switching and network reconfiguration^36^. Finally, we correlated the sensorimotor and neuropsychological performance of ataxic subjects, measured through clinical and neuropsychological assessments, with anatomo-physiological parameters and used unsupervised machine learning to verify whether the clinical groups are predicted by the underlying multiparametric space. This novel approach revealed profound differences in SMN and VAN organization and function between JS and SP, found commonalities at network level within SP despite the heterogeneous etiopathogenesis, and identified the basis for functional compensation in JS.

## Methods

### Subjects

A total of 16 patients with cerebellar ataxia were recruited at the Istituto Neurologico “Carlo Besta”, comprising 8 subjects with JS (3 females 21±6 years) and 8 with SP (3 females, 20±5 years) were enrolled. The patient group included those with a diagnosis of genetically confirmed paediatric cerebellar ataxic syndrome and cerebellar abnormalities (vermal/hemispheric/global hypoplasia/atrophy) on brain MRI. Additionally, 11 healthy controls (HC; 5 females, 21±2 years) were included. Exclusion criteria for all participants included contraindications to MRI, history of other neurological disorders and psychiatric illness. Controls were further excluded in the presence of cardiovascular or cerebrovascular risk factors, history of brain injury, psychiatric disorders, panic disorder, psychoactive treatment, recent headache within the week before imaging, or a family history of cluster headache. All participants were required to avoid any drugs for at least 24 hours prior to the MRI scan. The study was approved by the local ethics committee of the Istituto Neurologico “Carlo Besta” and conducted in accordance with the Declaration of Helsinki. Written informed consent was obtained from all participants or their legal guardians if under the age of 18 years old.

### MRI acquisition

MR images were acquired using a 3T Philips Achieva MRI scanner. The protocol was standardized across the Italian Neuroscience and Neurorehabilitation network^37^. The protocol included a high-resolution T1-weigthed (3DT1) image, a multi-shell diffusion weighted (DW) scan and a resting-state functional MRI (rs-fMRI). The 3DT1was acquired with a 3D fast field echo (FFE) sequence with repetition time (TR)=8.2 ms, echo time (TE)=3.8 ms, inversion time (TI)=855 ms, 240 sagittal slices and a spatial resolution of 1×1×1 mm^3^. The DW images were acquired with a spin-echo (SE) - echo planar imaging (EPI) sequence with TR=8400 ms, TE=85 ms, 32 non-collinear DW directions per b-value, b-value=1000/2000 s/mm^2^ and 7 no-DW images (for a totality of 71 volumes), 60 axial slices and a spatial resolution of 2.5×2.5×2.5 mm^3^. The rs-fMRI protocol was acquired with a gradient echo (GE) – EPI sequence with TR=2400 ms, TE=30 ms, 200 volumes, 40 axial slices and a spatial resolution of 3×3×3 mm^3^ with a gap=0.5 mm, resulting in a total acquisition time of 8 minutes.

### Clinical and neuropsychological assessment

A comprehensive battery of clinical and neuropsychological evaluations was performed on a subset of 19 subjects (8 HC, 5 JS, and 6 SP). Several cognitive domains were evaluated testing intelligence, visuomotor, executive, attentive, and emotional/social functions (see Supplementary for a detailed description). The battery included the Wechsler Adult Intelligence Scale – Fourth Edition (WAIS-IV)^38^, the Cerebellar Cognitive Affective Syndrome (CCAS)^39,40^ scale, the Rey–Osterrieth Complex Figure (ROCF)^41^, the Tower of London (ToL)^42^, the Continuous Performance Test – Third Edition (CPT3)^43^, the Adult Behaviour Checklist (ABCL)^44^, and the Social Responsiveness Scale – Second Edition (SRS2)^45^. Cerebellar and ataxia-related dysfunctions were additionally assessed in patients only using the Scale for the Assessment and Rating of Ataxia (SARA)^46^.

### Data preparation

#### DW and rs-fMRI preprocessing

The preprocessing of DW data included the Marchenko-Pastur principal component analysis (MP-PCA) for denoising, and the correction for susceptibility distortions, eddy currents, and subject motion. The preprocessing of rs-fMRI data included MP-PCA denoising, slice-timing correction, realignment, polynomial detrending, nuisance regression with 24 motion parameters and cerebrospinal fluid (CSF) temporal signal^47^, and temporal band-pass filtering (0.008-0.09 Hz).

#### Creation of the atlas

An *ad hoc* atlas consisting of 146 regions of interest (ROIs) was created as follows: 100 cerebral cortical ROIs were extracted from the Schaefer atlas^48^, 15 cerebral subcortical ROIs were extracted from the FMRIB’s Integrated Registration and Segmentation Tool (FIRST) atlas^49^, 25 cerebellar cortical ROIs were extracted from the CerebNet^50^ and 6 deep cerebellar nuclei ROIs were extracted from the spatially unbiased atlas template of the cerebellum and brainstem (SUIT) atlas^51,52^. CerebNet was chosen for cerebellar cortical segmentation instead of SUIT, as it is a validated deep learning-based method that offers higher segmentation accuracy, especially in patients with ataxia and hypoplastic cerebellar volumes. Moreover, CerebNet performs the segmentation directly in the subject’s space, whereas SUIT relies on non-linear atlas registration from MNI to subject space. The cerebellar nuclei were instead identified using SUIT as they are not included in CerebNet.

All 146 ROIs were exclusively assigned to one of the seven canonical functional networks (somatomotor, dorsal attention, ventral attention, visual, limbic, frontoparietal, and default mode)^48^.The Schaefer atlas already provides the mapping of the cerebral cortical ROIs to the seven networks. The cerebellar cortical regions were mapped by overlaying the CerebNet atlas to the Buckner functional atlas^53^ and assigning each ROI to the network with the highest membership probability, while the subcortical ROIs were assigned to the network based on literature^54^. Focusing on motor and attentive involvement in ataxia, this study included only two of the seven canonical functional networks, i.e., the SMN and the VAN, for subsequent analysis. Supplementary Table 1 lists the ROIs included in SMN and VAN.

#### Creation of structural and functional connectivity matrices

For each subject, structural connectivity (SC) and functional connectivity (FC) matrices were computed for each network, as detailed below.

Different brain tissues, i.e., cortical grey matter (GM), white matter, deep GM and CSF, were segmented from 3DT1 images. After applying a rigid registration between 3DT1 space and DW space, the tissue maps were applied as anatomical priors for anatomically constrained tractography^55^. The fiber orientation distributions were estimated using the multi-shell multi-tissue constrained spherical deconvolution and whole brain tractography was performed using probabilistic streamline tractography^56^ with 30 million streamlines. The tracts belonging to the cerebello-thalamo-cortical and the cortico-ponto-cerebellar tracts were anatomically curated to remove false positive streamlines^57^. Whole brain tractography was combined with SMN and VAN atlases to extract streamlines (edges) between brain ROIs (nodes), resulting in two SC matrices per network: a distance matrix (length-SC), representing the length of tracts connecting each pair of nodes, and a weight matrix (weight-SC), in which connection strengths are given by the normalized number of streamlines (divided by the maximum value).

Further, static FC and dynamic functional connectivity (FCD) matrices were computed for SMN and VAN. The FC matrix was calculated by extracting the average BOLD signal from each node within the network and calculating the Pearson correlation coefficients (PCC) between all pairs of ROI time-series. The resulting correlation values were transformed using a thresholded Fisher’s z-transformation^58^. FCD matrix, which instead captures the temporal evolution of functional interactions across brain regions during resting state, was computed by applying a sliding time window of 40 seconds to the BOLD time-series, incrementally shifted by 1 TR, resulting in a series of FC matrices at successive time points. Each matrix was vectorized, and pairwise correlations between these vectors were calculated to generate the final FCD matrix^59^ (178×178 in our case).

### Volume estimation

For each subject, the 3DT1 image was used to calculate normalized volumetric measures at multiple levels of analysis. First, the total volume of each network (SMN and VAN) was obtained by summing the volumes of all involved regions (*Vi*, calculated as the number of voxels belonging to region *i* multiplied by the voxel volume) and then divided by the intracranial volume (ICV) of the subject:

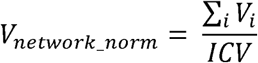

To evaluate the specific contribution of the cerebellum, the total normalized volume of all the cerebellar regions within each network was calculated in the same way. In addition, normalized volumes of each individual region within the networks were calculated to perform region-specific analyses.

To capture cerebellar volumes beyond the two networks, normalized volumes were computed for all cerebellar regions not assigned to either SMN or VAN. The cerebellar regional volumes were further summed to compute the total normalized volumes for lobules and vermis, providing a comprehensive anatomical characterization of cerebellar involvement.

### Graph theory

Graph theory is a mathematical framework used to model and quantify the topological properties of complex networks. In this context, brain networks are represented as graphs, where each region is modelled as a node (or vertex), and each anatomical or functional connection between regions is represented as an edge. The following network measures, grouped by type, were calculated from weight-SC and static FC matrices^60^ using the Brain Connectivity Toolbox (brain-connectivity-toolbox.net):

- Topological index: *mean edge weight* defined as the average of all edges, including zero-weighted (absent) edges;
- Centrality indices: *density* calculated as the ratio of existing edges and the total number of possible edges; *strength* calculated first by computing the sum of the edges for each node and then averaging these values across all nodes; *betweenness centrality* calculated first for each node as the proportion of times a node lies in the shortest path between all other pairs of nodes and then averaging these values across all nodes; and *core nodes,* derived from the core-periphery structure, were defined as the number of nodes assigned to the densely connected core (central hubs) rather than to the sparsely connected periphery in a binary partition of the network obtained by maximizing a core-ness statistic^61,62^;
- Integration index: *global efficiency* calculated as the average inverted shortest path length between each pair of nodes in the network;
- Segregation index: *clustering coefficient* defined as the average prevalence of strongly interconnected triangles around each node, measured as the proportion of node’s neighbour that are also neighbours of each other^63^.

In addition, the contribution of cerebellar nodes was considered to understand how much the cerebellum affected the topology of the network.

### Virtual brain modelling

The Virtual Brain (TVB) framework^30^ was used to simulate the neural dynamics of brain networks (i.e., SMN and VAN), assigning the Wong-Wang neural mass model to simulate the neuronal activity of each GM region (node)^32^. This model captures local microcircuit activity through two interacting populations of excitatory and inhibitory neurons, coupled via NMDA and GABA receptor-mediated synapses. Within TVB, each brain region was represented by this neural mass model, while inter-regional connectivity was constrained by subject-specific SC matrices reflecting connection strengths and distances. Model parameters were fixed according to values reported by Deco et al.^32^, except for four key parameters (global coupling and three synaptic parameters) optimized to assess subject-specific excitation/inhibition balance:

- Global coupling (*G)*: a scalar parameter that globally scales the strength of long-range excitatory connections between brain regions;
- Inhibitory synaptic coupling (J_i_): a local parameter that represents the strength of the inhibitory feedback within each brain region, mediated by GABAergic synapses;
- Excitatory synaptic coupling (J_NMDA_): a local parameter that represents the strength of the excitatory input to neuronal populations, mediated by NMDA receptors (glutamatergic synapses);
- Recurrent excitation (w^+^): a local parameter that represents the strength of self-excitation mediated by NMDA receptors within local excitatory populations.

For each subject, optimal parameters were determined using a grid search, in which large-scale neural dynamics were simulated for 8 minutes (matching the duration of rs-fMRI acquisition) for each combination of parameters within the predefined search space^64^. In detail, the Balloon-Windkessel hemodynamic model^65^ was applied to the simulated synaptic activity to generate resting-state BOLD time-series, which were used to compute simulated FC and FCD matrices.

Then, the PCC between empirical and simulated FC and the Kolmogorov-Smirnov (KS) distance between empirical and simulated FCD were calculated. Optimal parameters were determined by minimizing a cost function designed to maximize agreement between empirical and simulated FC and FCD^66^:

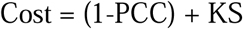

All simulations were performed on the Leonardo and Galileo high performance computing clusters at CINECA.

### Statistical analysis and machine learning

All features extracted from MRI data, including volumetric measures, graph theory indices, and TVB parameters, are collectively referred to as MRI derived parameters. Figure 1 illustrates an overview of how MRI derived parameters were computed. These parameters were subsequently used as predictor variables in the statistical and machine learning analyses described below.

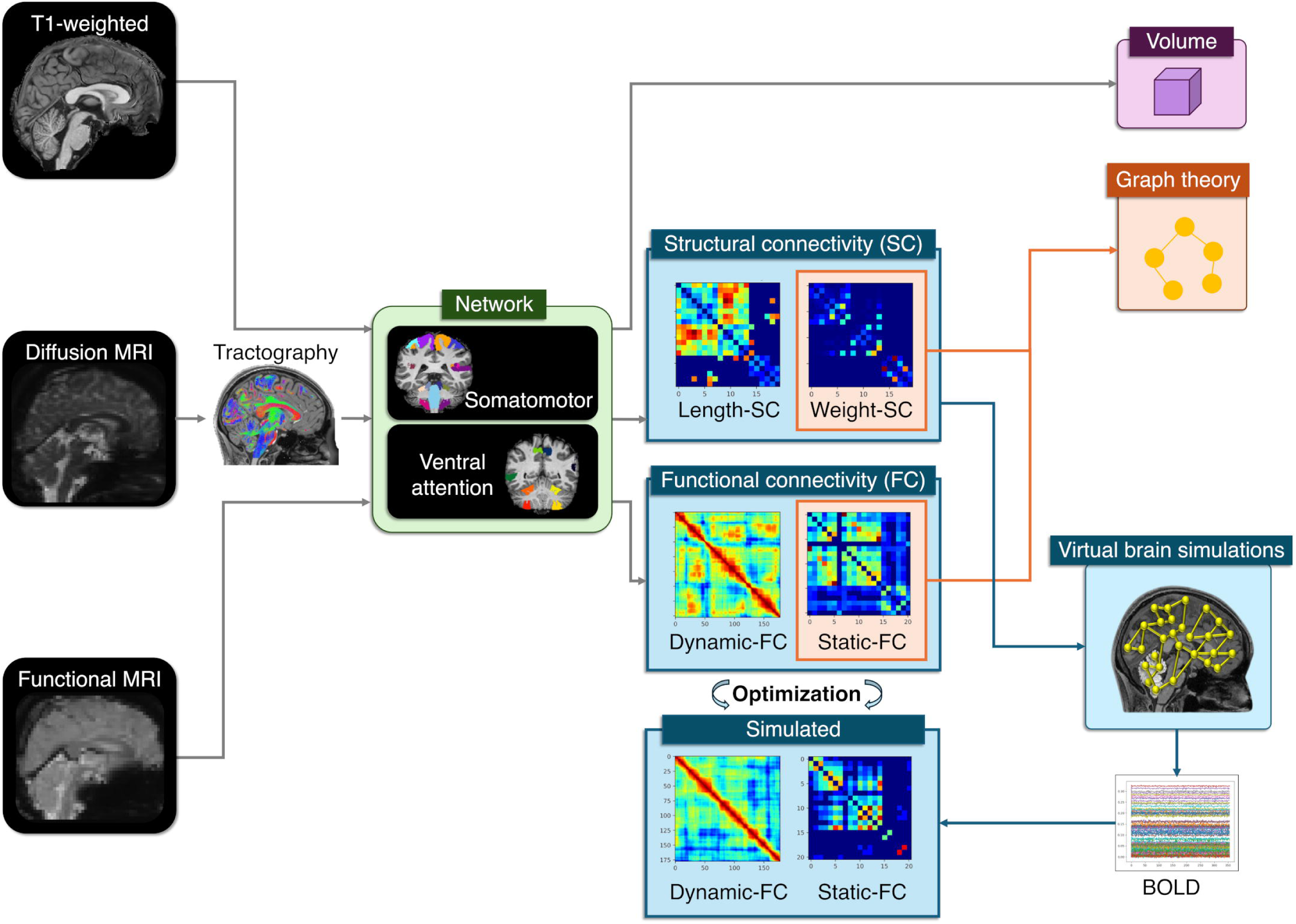

#### Assessment of group differences

All MRI derived parameters for the SMN and VAN were compared between groups to assess whether significant differences existed between patients and controls. The groups were matched for age and sex. The statistical test was chosen based on data distribution: when normally distributed, a one-way ANOVA followed by Tukey’s post hoc test was used, otherwise a Kruskal-Wallis test followed by Dunn’s post hoc test was applied. The statistical threshold was always set at p=0.05. The same statistical analysis was applied to clinical and neuropsychological scores, including ABCL, CCAS, CPT3, REY, RTT, SARA, SRS_SCI, Tol, and WAIS-IV measures, to assess group differences. For the following analyses, only MRI derived parameters that were significantly different between groups in at least one of the networks were considered.

#### Assessment of the relation between clinical/neuropsychological scores with MRI derived parameters

To assess the ability of MRI derived parameters to predict clinical and neuropsychological scores, a stepwise linear regression for each score was performed iteratively eliminating variables that did not contribute significantly to the model. Clinical or neuropsychological scores were used as dependent variables, while all MRI derived parameters were included as independent variables. To reduce the risk of overfitting, only network level volumetric measures were considered, whereas cerebellar and individual region volumes were excluded.

The best robust linear regression models were selected based on the adjusted coefficient of determination (R²-adj), set to be greater than 0.6, and normalized root mean square error (NRMSE), set to be less than the median.

#### Clustering

To assess whether selected combinations of MRI derived parameters can identify biologically relevant groups within each network, the correspondence between unsupervised clustering and the clinical group membership (i.e. HC, JS and SP) was explored. In addition to the total network volume, we also included volumes of single regions that differed significantly between groups.

For each network, selected MRI-derived parameters were normalized to zero mean and unit standard deviation before performing principal component analysis (PCA). In this way, data dimensionality was reduced preserving the components with the highest variance, i.e., the first two principal components (PC1 and PC2). K-means clustering^67^ was applied to the space reduced by PCA with 100 random initializations to ensure robust convergence. To choose the optimal number of clusters (k), we used the gap statistic method, which compares the within-cluster dispersion of the observed data to that expected under a randomly generated null distribution. The method was applied to evaluate cluster numbers from 2 to 5, using 1000 bootstrap samples. The optimal k was defined as the smallest number of clusters for which the gap statistic satisfied the criterion Gap(k) ≥ Gap(k+1) − SE(k+1), where SE represents the standard error of the gap statistic^68^.

## Data Availability Statement

The structural and functional connectivity matrices are publicly available on Zenodo at: https://doi.org/10.5281/zenodo.18862694.

## Results

A comprehensive analysis of morphometric, topological, and dynamical network parameters (Figure 1) was carried out on 8 patients diagnosed with JS, 8 patients diagnosed with SP, and 11 HC. The SP group included patients with spinocerebellar ataxia (SCA13^18^, SCA29^20^), coenzyme Q deficiency^21^, autosomal recessive spastic ataxia of Charlevoix-Saguenay (ARSACS)^69^, peroxisomal disorders^70^, ITPR1-related ataxia^19^, congenital disorders of glycosylation type I (CDG-Ia)^71^, mitochondrial respiratory chain complex II deficiency^72^. Genetic information for patients with ataxia (the associated genes and sequence variances) is reported in Supplementary Table 2.

### Volumetric alterations

SP patients exhibited a widespread reduction in cerebellar volume compared to HC and JS patients. JS patients, in contrast, showed more localised cerebellar alterations (Figure 2), confirming classical neuropathological and neuroradiological findings^16^.

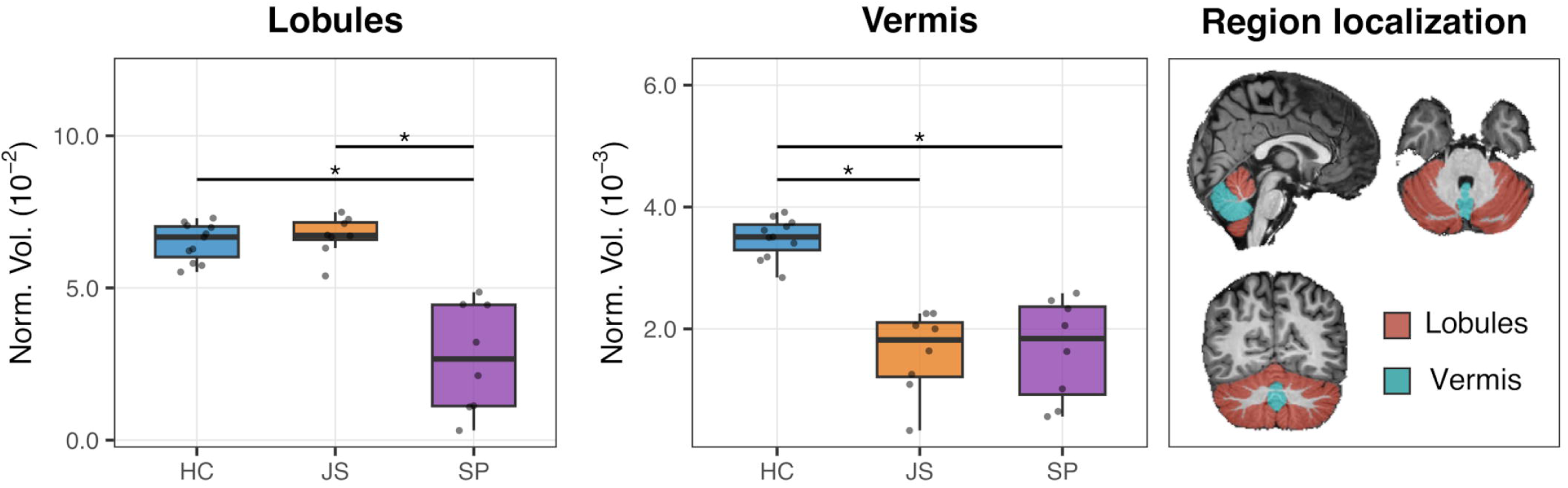

The cerebellar atrophy reflected into volume reduction in SMN and VAN (Figure 3). In SMN, SP patients exhibited a reduced volume compared to HC (p=0.008). When considering only cerebellar regions of SMN, SP patients showed a reduced volume compared both to HC and JS patients (Figure 3). In VAN, both JS (p=0.03) and SP patients (p=0.003) showed a reduced volume compared to HC. When considering only cerebellar regions of VAN, SP patients showed a reduced volume compared to both HC and JS (p<0.0001), whereas no significant difference was observed between JS and HC (Figure 3). Therefore, network volume reduction in SMN and VAN was more extensive in SP than JS and was almost entirely explained by changes occurring in the cerebellum.

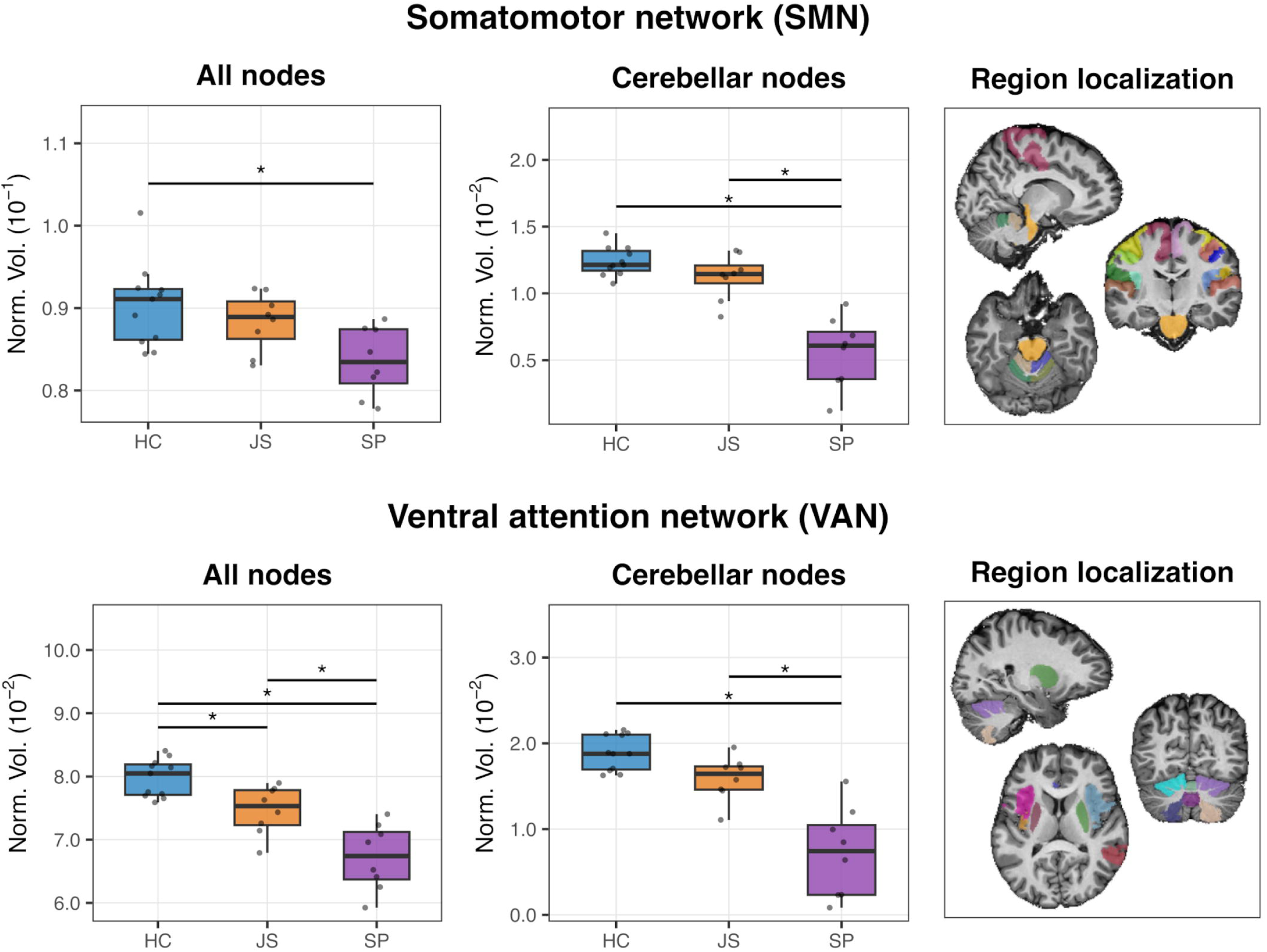

Examining individual regions, statistically significant group differences were observed exclusively in cerebellar regions (Figure 4). In SMN, all the six cerebellar regions showed a reduced volume in SP patients compared to HC and JS patients. In addition, bilateral lobules I-IV were reduced in JS patients compared to HC. In VAN, all six cerebellar regions were reduced in SP patients compared to HC. Furthermore, bilateral lobules V and bilateral lobules VIIIa were reduced in SP patients compared to JS patients. Moreover, vermis VI, vermis VIII and right lobule VI were reduced in JS patients compared to HC (Figure 4). Therefore, network volume reductions were also detected in smaller regions of the cerebellum.

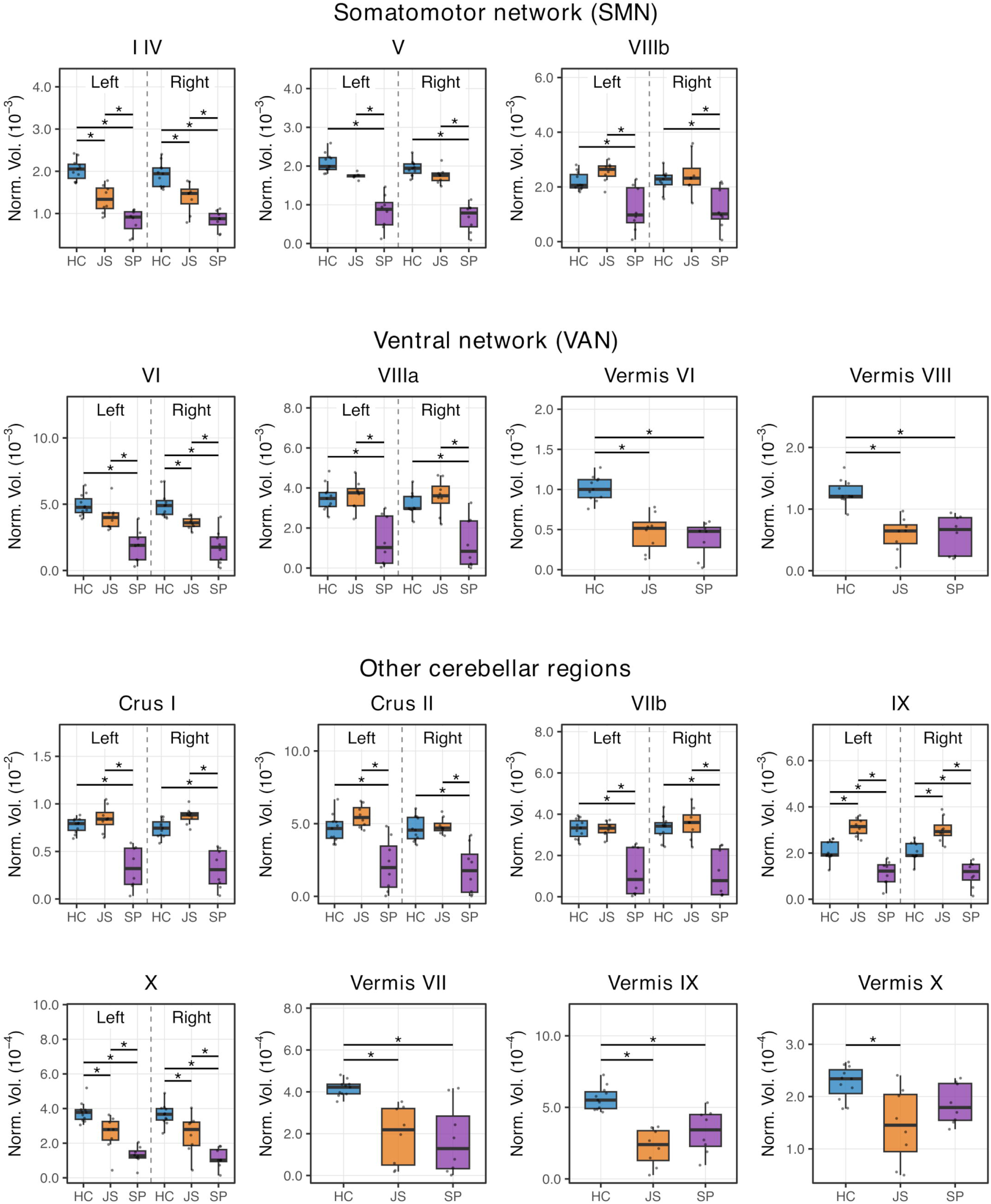

Finally, when considering all cerebellar regions individually, SP patients showed a volume reduction compared with HC in all regions (p≤0.001) except vermis X. JS patients showed a volume reduction compared with HC in all vermis regions and in bilateral lobule X (p≤0.001) but a volume increase in bilateral lobule IX (Figure 4).

### Topological alterations

Graph theory metrics, which were used to quantify changes in network topological organization, revealed significant group differences in weight-SC matrices for both the SMN and VAN, and in FC matrices for the VAN only (Figure 5).

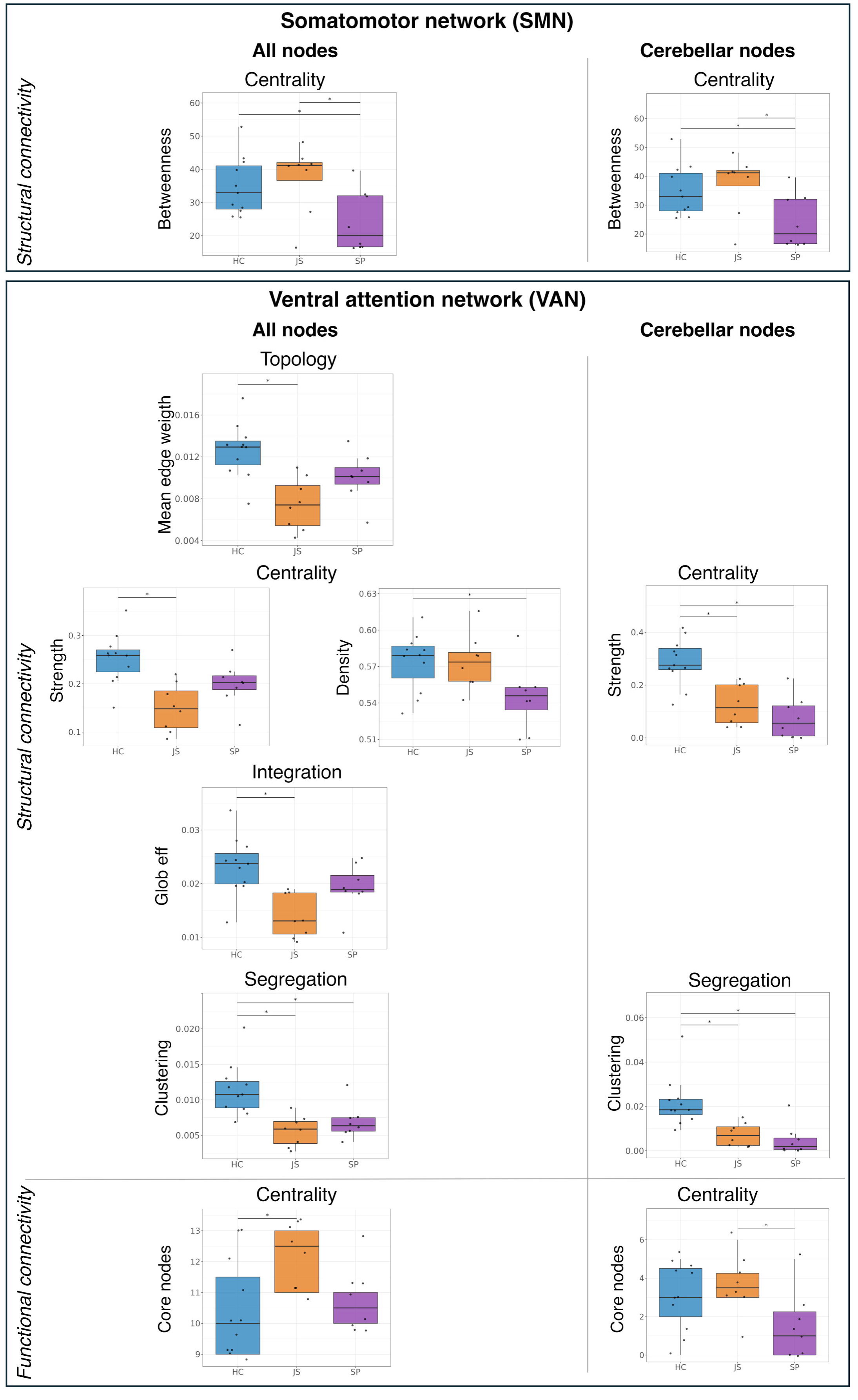

In SMN, SP patients showed reduced mean betweenness centrality in weight-SC matrices compared to both HC (p=0.034) and JS patients (p=0.018), meaning that SMN regions in SP were structurally less central. The same result was confirmed when considering only cerebellar nodes belonging to SMN.

In VAN, both JS and SP patients showed altered structural organization compared to HC. Specifically, JS patients showed reduced mean edge weight (p=0.0004), strength (p=0.0004), and global efficiency compared to HC (p=0.0008), reflecting a widespread weakening of structural connections. On the other hand, SP patients showed reduced density (p=0.038), indicating that the number of structural connections was reduced but the remaining connections maintained their strength. In addition, both JS (p=0.0008) and SP patients (p<0.008) showed reduced clustering coefficient in the weight-SC matrices, indicating a loss of network structural segregation. Similar results were observed when considering only the cerebellar nodes in VAN. Indeed, both JS (p=0.007) and SP patients (p=0.0005) showed a reduced clustering coefficient and both JS (p=0.0009) and SP patients (p=0.00003) showed reduced strength compared to HC.

Despite structural alterations in VAN, JS patients showed an increased number of functional core nodes (derived from FC matrix) compared to HC (p=0.01). When considering only the cerebellar nodes in VAN, SP patients showed a decrease in the number of functional core nodes compared to JS (p=0.049).

### Alterations in network dynamics

TVB analysis gave information about the connectivity between nodes and their excitation/inhibition balance^64,73,74^. TVB parameters revealed group difference in both SMN and VAN networks (Figure 6). In SMN, SP patients showed higher Ji values compared to JS patients (p=0.02). In VAN JS patients showed higher G compared to HC (p=0.01).

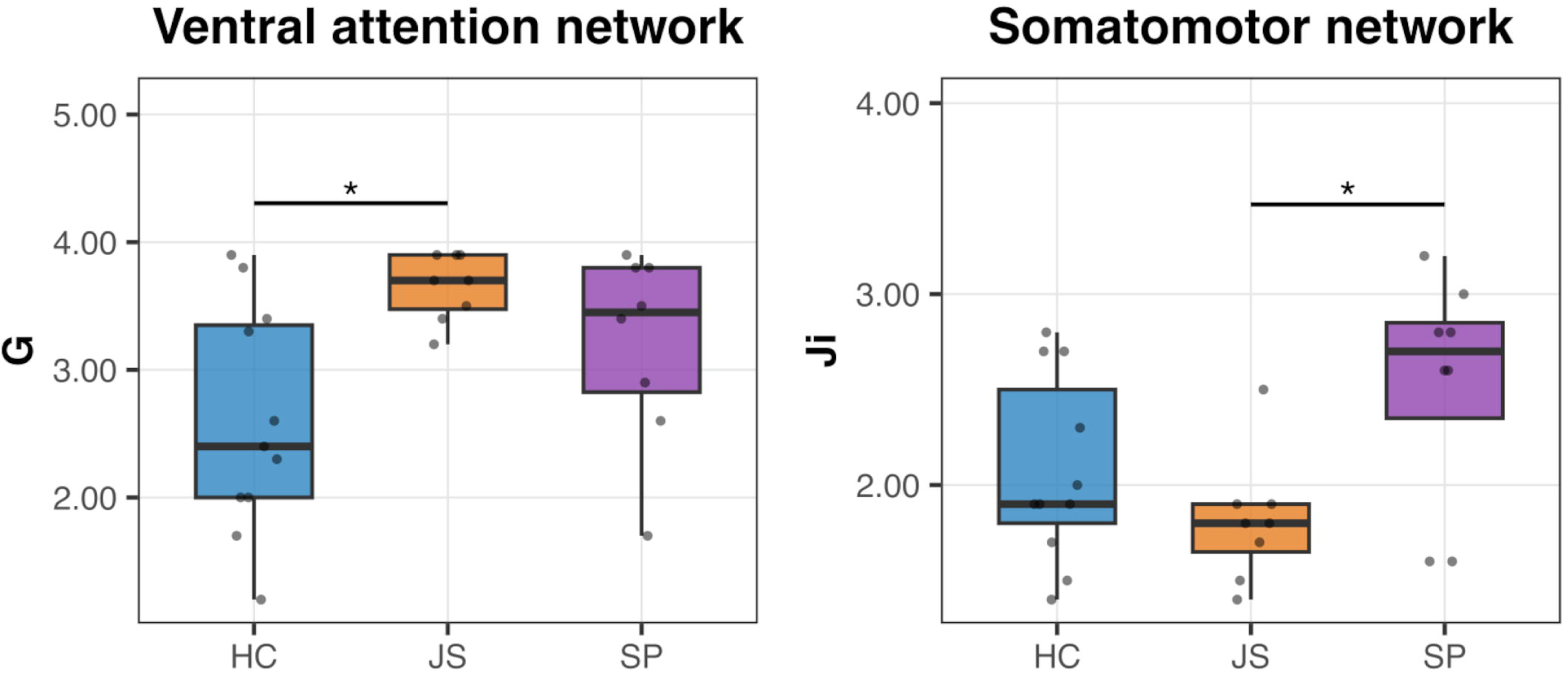

### Correlation of clinical and neuropsychological scores with multiple network parameters

Group differences in clinical and neuropsychological scores between SP, JS and HC are reported in Supplementary Table 3. More interestingly, the subjective multiparametric space was correlated with clinical and neuropsychological scores. A stepwise regression analysis was used to identified MRI-derived network parameters that reliably predict clinical and neuropsychological scores. Only the best performing regression models are summarized in a heatmap (Figure 7A), and two representative regression examples are shown as scatter plots (Figure 7B), while the scatter plots for remaining significant regression models are reported in Supplementary Figure 1. Notably, two regression models within SMN and fourteen within VAN demonstrated strong predictive power (R^2^-adj>0.6 and NRMSE<median). Therefore, morphological, topological, and dynamical network parameters revealed a strong predictive power for the sensorimotor and cogntive consequences of ataxia.

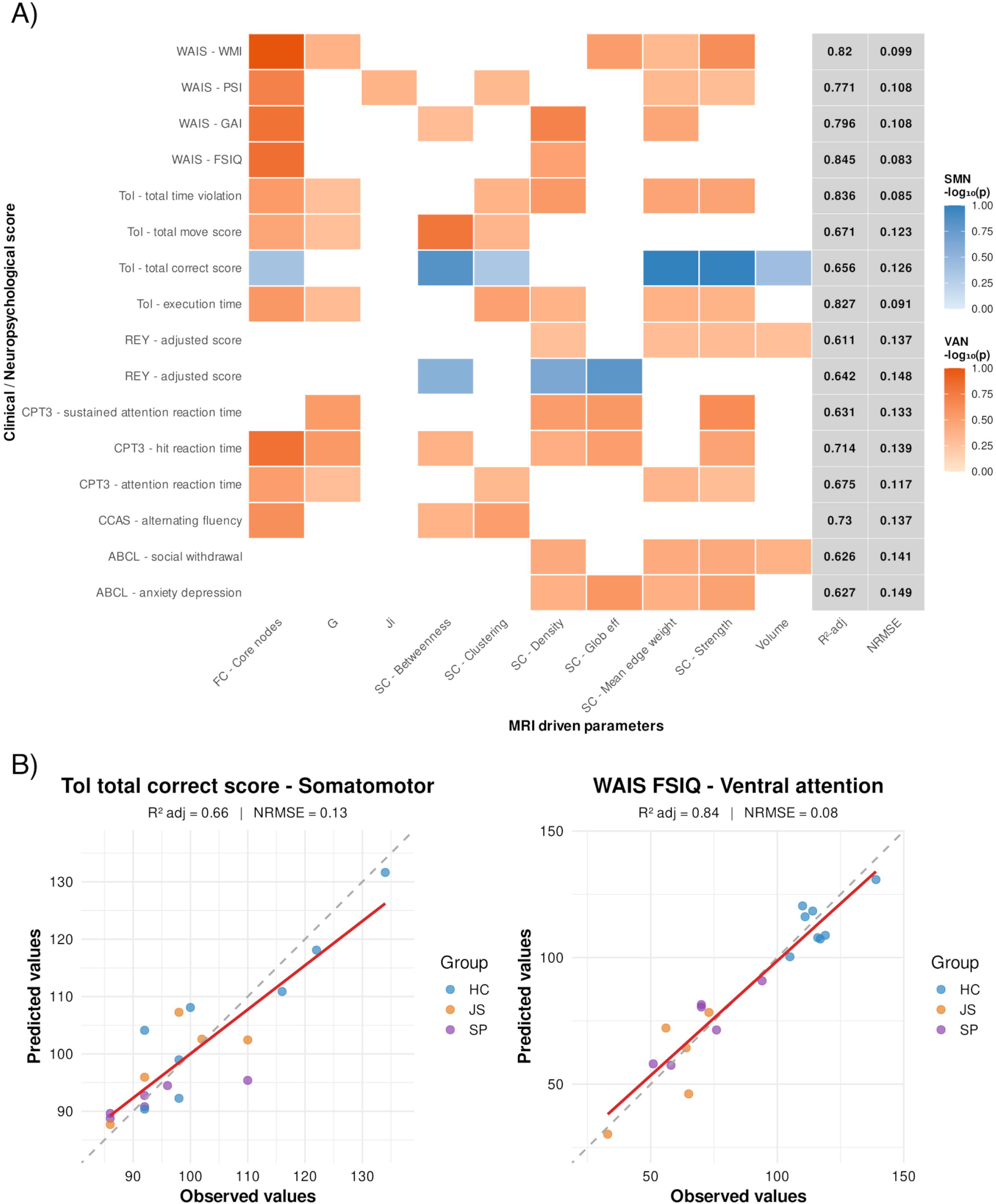

### Clustering

For each network, PCA was applied to MRI-derived parameters to reduce dimensionality and identify major patterns of variance. PCA was followed by k-means clustering to identify biologically meaningful subgroups (Figure 8).

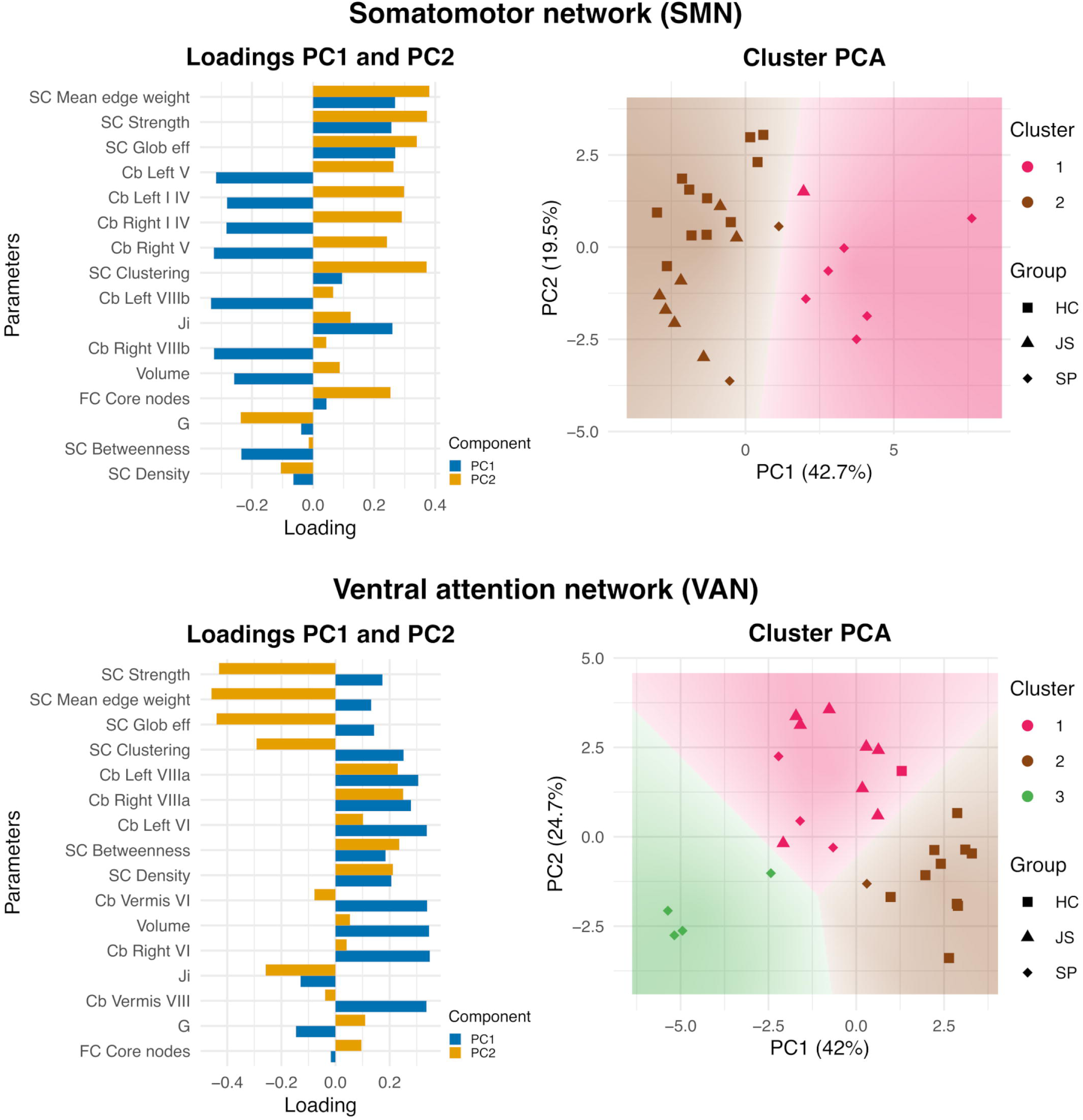

In SMN, the first two principal components (PC1 and PC2) explained a total of 62.2% of the variance in the MRI-derived parameters, with PC1 accounting for 42.7% and PC2 for 19.5%. The MRI-derived parameters with the highest loadings were volumes, such as volume of left cerebellar lobule I-IV and V, and metrics derived with graph theory, such as SC mean edge weight, SC strength, and SC global efficiency. K-means clustering assigned subjects to two clusters. Cluster 2 included all HC (11/11), 7 out of 8 JS patients, and 2 SP patient, whereas cluster 1 consisted of 6 out of 8 SP patients and 1 JS patient.

In VAN, the first two principal components explained a total of 66.7% of the variance in the MRI-derived parameters, with PC1 accounting for 42% and PC2 for 24.7%. The MRI-derived parameters with the highest loadings for PC1 and PC2 were volumes, such as volume of left cerebellar lobule VIIIa, and metrics derived with graph theory, such as SC strength, SC mean edge weight, SC global efficiency and SC clustering coefficient. K-means clustering assigned subjects to three clusters. Cluster 1 included all JS subjects (8/8), 3 SP subjects and 1 HC subject. Cluster 2 included mostly HC subjects (10/11) and 1 SP patients while cluster 3 included only 4 SP patients.

## Discussion

Beyond cerebellar atrophy^75^, brain circuit topology and dynamics reveal strong alterations in signal elaboration and communication between several cerebral regions and the cerebellum, with patterns of change that differentiate JS from SP, as well as these two forms of ataxia from HC. Using complementary morphometric, topological, and dynamic network measures to construct subject- specific brain digital twins, we characterised large-scale network alterations associated with cerebellar pathology and individual motor and cognitive performance.

Specific alterations in the SMN and in the VAN strongly correlate with the motor and affective-cognitive performance of subjects, so that clinical and neuropsychological scores are predicted by anatomo-physiological network parameters. Indeed clinical and neuropsychological scores were predicted by parameters of their respective functional networks: for instance, the Tower of London was associated with SMN, while the Wechsler Adult Intelligence Scale (WAIS) was associated with VAN. As a validation of the informative capacity of these parameters, PCA and clustering demonstrated that MRI-derived parameters are sufficient to discriminate the JS, SP and HC groups.

### Cerebellar atrophy patterns

Volumetric analysis revealed distinct patterns of cerebellar involvement in SP and JS patients compared to HC. SP patients were characterized by widespread cerebellar atrophy affecting both hemispheric lobules and vermis, whereas JS patients showed more selective cerebellar involvement, predominantly affecting the vermis. Surprisingly, lobule IX (left and right) showed increased volume in JS patients compared to HC. Lobule IX is recognized for its role in social cognition, action sequencing, and higher order cognitive functions^76,77^. Interestingly, therapies such as cognitive orientation to daily occupational performance in children with developmental coordination disorder have been shown to increase grey matter volume in lobule IX and Crus I, at the same time improving motor performance^78^. Therefore, our finding may reflect a compensatory mechanism, potentially supporting functional adaptation in response to vermis hypoplasia or other cerebellar deficits typical of JS.

### The somatomotor network and motor deficits

SP patients showed a widespread cerebellar atrophy, affecting both hemispheres and the vermis, that brought about a reduced SMN volume compared to HC. Moreover, SP patients showed a reduced structural betweenness centrality (compared to HC), implying that network nodes cannot efficiently transfer information across the network. Taken together, these results show that the primary cerebellar damage secondarily impacts the entire network. Moreover, the local inhibitory feedback (Ji) mediated by GABAergic synapses was higher in SP than JS, reducing the excitation/inhibition balance. This could concur to reduce information transfer through the network together with the widespread network atrophy and reduced centrality. In contrast, JS patients showed only a selective reduction of the anterior vermis compared to HC but not significant changes in SMN parameters. Overall, these findings indicate that SMN abnormalities are more pronounced in SP than JS explaining the more severe sensorimotor deficit revealed by SARA scores (Supplementary Table 3). Furthermore, both SP and JS performed less efficiently than HC on the Rey Complex Figure Test, and the correlation with MRI-derived SMN parameters confirms the central role of sensory-motor functions, specifically visuomotor coordination and fine motor control^79^, alongside executive strategic planning^80^. Interestingly, unsupervised clustering revealed that SP patients share a common pattern of SMN parameters (75% were into the same cluster together with only one JS) despite the heterogeneity of etiopathogenetic factors.

### The ventral attention network and affective-cognitive deficits

In JS patients, cerebellar atrophy brought about a reduced VAN volume compared to HC. JS patients also featured relevant topological and dynamical changes (compared to HC): (i) a decreased mean edge weight and global efficiency, implying weaker connections and reduced network integration; (ii) a reduced strength and cluster coefficient, indicating weaker connection weights and decreased segregation; (iii) an increased number of functional core nodes, suggesting an increased information trafficking within the network; (iv) an increased global coupling, enhancing connectivity synchrony in the network. While the reduced integration and segregation of information reflect structural alterations, increase of functional core nodes and of their global coupling may be compensatory to preserve cognition and attention. These signs of compensation, observed only in JS subjects, may reflect the greater developmental plasticity associated with the congenital nature of JS, which could support more effective functional reorganization. Interestingly, when considering functional connectivity only for cerebellar nodes, core nodes and global coupling are no longer significantly increased, akin with the concept that the structural damage is primarily localized within the cerebellum, while the functional compensation involves extracerebellar nodes. In contrast, SP patients (compared to HC) showed structural alterations only in clustering coefficient, indicating a decrease of the network segregation, and a reduced node density, reflecting fewer structural connections. Overall, these findings indicate that VAN changes show signs of compensation in JS but not in SP. Interestingly, unsupervised clustering revealed that JS patients share a common pattern of VAN parameters (100% were into the same cluster together with only one HC and three SP).

### Cerebellar alterations and the physiopathology of ataxia

But how are the cerebellar alterations of JS and SP related to the physiopathology of ataxia? Theory predicts that conditions altering cerebellar maturation and functioning can impair its core algorithm (the so-called *universal cerebellar transform*)^81,82,25,83^ leading to motor and cognitive dysmetria and causing ataxia of movement and thought^84,85,86^. A more subtle question is how cerebellar alterations differentially impact JS and SP. Are there distinctive changes depending on subject and pathology or, rather, the breakdown of the cerebellar input-output function is non-specific, so that the impact reflects the connectivity of lesioned areas with large-scale brain circuits (the so-called *meta-level hypothesis)*^82,87^? Our results support the latter case. Indeed, the SMN of SP patients shows a common pattern of large-scale circuit alterations despite the many diverse etiopathogenetic causes (our sample includes alterations of K^+^ ionic channels^18^, calcium metabolism^19,88^, the cellular respiratory pathway^21^ the endocytic pathway^18^, the peroxidative system^18^, and protein glycosilation^18^). The VAN of JS patients also shows a common pattern of large-scale circuit alterations despite the numerous gene mutations involved^10,11,12^. Therefore, it is not the specific cellular alteration in the cerebellar microcircuit but rather the impairment of its input-output function in critical nodes of large-scale networks that explains the differential involvement of SMN and VAN in JS and SP.

### Study considerations

Despite the insight provided by these results, the study presents some limitations. First, the sample size is relatively small due to the rarity of these disorders, and this may limit the generalisation of our findings. Secondly, the definition of brain networks depends on the atlas, and this might influence the analyses (including volumetric, graph theory, and virtual brain simulations). Finally, virtual brain simulations were run using a standardized pipeline^64,73,74^ but future updates to the neural mass model are expected to improve the interpretation of pathological and compensatory mechanisms in the cerebellar network. These should include cerebellum specific mean field models^89^ and an anatomically refined structural connectivity^90.^

## Conclusions

Our analysis of brain circuit alterations allows to identify the neural underpinnings of two forms of ataxias, JS and SP. The degeneration of the cerebellum brings about structural, functional, and dynamical alterations of large-scale networks to which it is connected impairing the global capacity of the brain to carry out motor and affective-cognitive operations. Interestingly, the SMN turns out to be especially altered in SP and the VAN in JS. The nature of changes suggests that the primary damage coexists with compensatory mechanisms in JS. The connectivity of altered cerebellar areas into large-scale brain networks, along with extracerebellar compensatory mechanisms, determine the different balance and gravity of motor and affective-cognitive symptoms in JS and SP. Indeed, symptoms correlate with alterations of anatomo-physiological parameters in large-scale networks rather than with the etiopathogenetic (genetic or molecular) causes. To further understand the underlying mechanisms of damage and compensation, specific neurophysiological and neuropathological studies are warranted, combining electrophysiological recordings with neural network simulations^24,91^. This approach will allow experimental data to inform models, enabling the investigation of how changes in brain networks affect function and how the system self-repairs and adapts. These new insights on the network mechanisms underlying different forms of cerebellar ataxia may be exploited to design potential therapeutic interventions to potentiate compensatory mechanisms, e.g., using patient’s brain digital twins to simulate the impact of targeted neuromodulation^29^. Focal neurostimulation (e.g., using transcranial magnetic stimulation or deep brain stimulation)^92^ could be used to modify the excitation/inhibition balance in specific nodes or networks and be combined with neurorehabilitation and pharmacotherapy to establish long-term changes in the cerebellar circuits of interest^93^. Thus, the capacity of detecting network changes in single patients may be exploited in the future for better profiling and for tailoring personalized therapeutic strategies for ataxias.

## Funding

M.G. and C.C. are supported by the Piano Nazionale di Ripresa e Resilienza (PNRR) within the “National Centre for HPC, Big Data and Quantum Computing” project (No. CN00000013, PNRR MUR-M4C2-Fund 1.4, Call “National Centers”, Law Decree n. 3138, 16 December 2021).

E.D. is supported by the National Recovery and Resilience Plan (NRRP), project IR00011–EBRAINS-Italy, funded by the European Union – NextGenerationEU (Project IR0000011, CUP B51E22000150006, “EBRAINS-Italy”), and by The Virtual Brain Twin Project, which has received funding from the European Union’s Horizon Europe Research and Innovation Programme under grant agreement No. 101137289.

F.P. and C.C. are supported by the European Union’s Horizon Europe Programme under Specific Grant Agreement No. 101147319 (EBRAINS 2.0 Project).

C.A.M.G.W.-K. is supported by #NEXTGENERATIONEU (NGEU) and funded by the Ministry of University and Research (MUR) through the National Recovery and Resilience Plan (NRRP), project MNESYS (PE0000006) – “A multiscale integrated approach to the study of the nervous system in health and disease” (DN. 1553, 11 October 2022).

Data collection at Fondazione IRCCS Istituto Neurologico C. Besta was supported by the Italian Ministry of Health (RC 2025) and Fondazione Pierfranco and Luisa Mariani for child neurology.

## Competing interests

The authors report no competing interests.

## Supplementary material

Supplementary material is available at *Brain* online.

## Notes

### Competing Interest Statement

The authors have declared no competing interest.

