## Supplementary for "Brain digital twins reveal network changes in congenital and slowly progressive cerebellar ataxias"

**Details of clinical and neuropsychological scores**

Below are the descriptions of all clinical and neuropsychological scales used in the study, with the specific outcome measures that were analysed.

1) Scale for the assessment and rating of ataxia (SARA)[43] is a clinical scale used to evaluate the severity of ataxia in terms of motor and functional impairment. Raw scores range from 0 (no ataxia) to 40 (maximum severity of ataxia).
2) Wechsler adult intelligence scale (WAIS)-IV[34] is a neuropsychological test used to assess different aspects of intelligence. Specifically, the following indices were considered:
   - Full Scale Intelligence Quotient (FSIQ) is the most representative estimate of overall intellectual functioning.
   - General Ability Index (GAI): provides an estimate of general intellectual ability considering the verbal comprehension and perceptual reasoning.
   - Working Memory Index (WMI): measures the ability to pay attention to verbally presented information, to manipulate it in immediate short-term memory and to formulate a response.
   - Processing Speed Index (PSI): assesses the speed of cognitive processes and responses.

   All composite indices were expressed as standard scores with mean of 100 and standard deviation of 15.
3) Cerebellar cognitive affective syndrome scale[35,36] (CCAS-s) is a brief clinical screen composed by 10 subtests designed to detect the cognitive and emotional deficits specific to cerebellar damage in addition to classic motor problems. All JS and SP subjects showed a definite CCAS (3 or more failed subtests), except one SP who failed only 2 subtests (probable CCAS). The raw scores obtained in the following executive function and affect subtests were included in statistal analysis of correlation between neuropsychological and MRI derived parameters:
   - Alternating fluency/Category Switching: tests cognitive flexibility. The patient has to generate words alternately from two categories (e.g., naming a fruit, then a piece of furniture), measuring the ability to shift between criteria without rigidity.
   - Digit Span Backward: the clinician says “a”, and the patient must repeat “a” string of numbers, verbally presented by examinator, in reverse order. This is a classic measure of verbal working memory.
   - Go-NoGo: the patient must respond to specific stimuli (‘Go’) and inhibit responses to others (‘NoGo’). This test assesses the efficiency of motor inhibition and control of impulsivity responce.
   - Affectivity: the clinician evaluates through behavioral observation the presence of flat emotional response, disinhibition and emotional lability.
4) The Tower of London (ToL)[37] assesses executive functions, particularly planning, problem solving and inhibitory control. All scores were expressed as standard scores with mean of 100 and standard deviation of 15. The following scores were assessed:
   - Total correct score: total number of problems solved correctly within the minimum number of moves required.
   - Total move score: total number of moves made during the test.
   - Initiation time: time taken by the patient before making the first move.
   - Execution time: total time elapsed from the start of execution until the problem is completed, excluding the start time.

- Total time violation: number of times the patient fails to solve a problem within the predetermined maximum time.
- Rule Violation: number of times the patient violates the rules of the test.

5) Copy of Rey-Osterrieth complex figure test (ROCF) for visuomotor copying skills[38]. The raw is a raw score adjusted for age and education, that assesses accuracy, spatial organisation and proportionality.
6) Continuous performance test - third edition (CPT3)[40] assesses sustained attention, impulsivity and response inhibition. All variables were expressed as T-scores with mean of 50 and standard deviation of 10. The following scores were assessed:
   - Errors of omission: failure to respond to targeted stimuli.
   - Errors of commission: responses even when the stimulus did not require an action.
   - Errors of persistence: repetitive or excessive responses.
   - Hit reaction time variability: measures fluctuations in response speed.
   - Attention reaction time: assesses the speed of response to target stimuli.
   - Sustained attention reaction time: assesses changes in response time over the duration of the test.
   - Vigilance reaction time: measures the consistency of response to infrequent stimuli.
7) Adult Behaviour Checklist (ABCL)[41] is a standardised questionnaire used to assess emotional, behavioural and social functioning in adults. Specifically, the internalizing scales were assessed:
   - Anxiety/depression: measures symptoms of emotional distress, excessive worrying, sadness and nervousness.
   - Social withdrawal: assesses avoidance of social interactions, isolation and reduced interest in activities.
   - Somatic complaints: assesses physical symptoms without a clear medical cause, often related to psychological distress.
   - Total internalisation: global score combining anxiety/depression, social withdrawal and somatic complaints and reflecting the overall severity of internalisation problems.

   Subscales and Total Internalization score were expressed in T-score with mean of 50 and standard deviation of 10.
8) Social responsiveness scale-2 (SRS)[42] is a standardized questionnaire designed to assess social communication, social interaction, and associated restricted or repetitive behaviours. The social communication and interaction (SCI) scale is computed as part of the SRS. This score specifically evaluates social communication abilities, including understanding emotions, maintaining eye contact, and responding appropriately in conversations. It is expressed in T-score with mean of 50 and standard deviation of 10.

**Supplementary Table 1: Brain regions included in the somatomotor and ventral attention networks.**

The figure lists the cortical and subcortical regions comprising the somatomotor network (SMN) and the ventral attention network (VAN) analysed in this study.

| SOMATOMOTOR NETWORK (SMN) | VENTRAL ATTENTION NETWORK (VAN) |
|---|---|
| SomMot_1 (left and right) | SalVentAttn_ParOper_1 (left) |
| SomMot_2 (left and right) | SalVentAttn_FrOperIns_1 (left) |
| SomMot_3 (left and right) | SalVentAttn_FrOperIns_2 (left) |
| SomMot_4 (left and right) | SalVentAttn_PFCl_1 (left) |
| SomMot_5 (left and right) | SalVentAttn_Med_1 (left and right) |
| SomMot_6 (left and right) | SalVentAttn_Med_2 (left and right) |
| SomMot_7 (right) | SalVentAttn_Med_3 (left) |
| SomMot_8 (right) | Putamen (left and right)) |
| Brain-Stem | SalVentAttn_TempOccPar_1 (right) |
| Cb Lobule I-IV (left and right) | SalVentAttn_TempOccPar_2 (right) |
| Cb Lobule Left V (left and right) | SalVentAttn_FrOperIns_1 (right) |
| Cb Lobule VIIIb (Left and right) | Cb Lobule VI (left and right) |
| | Cb Lobule VIIIa (left and right) |
| | Cb_Vermis_VI |
| | Cb_Vermis_VIII |

**Supplementary Table 2. Genes and variants identified in patients with cerebellar ataxia.**

The table lists the genes associated with the clinical diagnosis, the mode of inheritance (autosomal dominant (AD) or recessive (AR)) and, the corresponding sequence variants identified in patients with Joubert syndrome and slowly progressive cerebellar ataxia.

| Group | Gene (OMIM) | AD/AR | Variants | |
|---|---|---|---|---|
| Joubert syndrome | AHI1 (*608894) | AR | c.1829G>C , p.R610P | c.2671C>T, p.R891X |
| | Not defined | | | |
| | TMEM67 (*107257) | AR | c.370G>A, p.E124K | c.1073C>T, p.P358L |
| | TMEM67 (*107257) | AR | c.370G>A, p.E124K | c.1073C>T, p.P358L |
| | CC2D2A (*612013) | AR | c.3289delG, p.Val1097fs | c.4667A>T, p.Asp1556Val |
| | CPLANE1 (* 614571) | AR | c.2747-161A>G; p.Gly916Gly_Ala917ins21 | Chr5:37238959-37239971 del |
| | ARMC9 (* 617612) | AR | c.1840_1853del, p.Leu614AlafsTer23 | c.991C>T; p.Arg331Cys |
| | CEP290 (*610142) | AR | c.2368-1G>A | c.5709+2T>G |
| Slowly progressive | KCN3C (* 176264) | AD | c.1268G>A, p.Arg423His | / |
| | ADCK3 (* 606980) | AR | c.547C>T, p.Gln183* | c.1042C>T,p.Arg348* |
| | ITPR1 (* 147265) | AD | c.805C>T, p.Arg269Trp | / |
| | SACS (*604490) | AR | | |
| | PEX12 * 601758 | AR | c.1015T>A, p.Cys339Ser | c.969_970delGT , p.Phe324LeufsTer29 |
| | PMM2 (*601785) | AR | | |
| | ITPR1 (* 147265) | AD | c.7471G>C, p.Gly2491Arg | / |

**Supplementary Table 3: Clinical and neuropsychological scores across groups.**

This table reports clinical and neuropsychological scores for three groups: HC = healthy controls, JS = Joubert syndrome, and SP = slowly progressive. Values are presented as mean (standard deviation) or Median (range), based on normal or non-normal distribution of test scores. The p-values indicate the results of statistical tests comparing all three groups (ANOVA* or Kruskal-Wallis§). Post-hoc pairwise comparisons are provided as HC vs JS, HC vs SP, and JS vs SP.

| | **HC**<br>**N 8** | **JS**<br>**N 5** | **SP**<br>**N 6** | **p-value** | **HC vs JS** | **HC vs SP** | **JS vs SP** |
|---|---|---|---|---|---|---|---|
| **SARA** | | 9.8 (5.12) | 16.5 (3.27) | 0.0414* | | | |
| **WAIS-IV** | | | | | | | |
| **FSIQ** | 116.4 (10.2) | 58.2 (15.3) | 69.8 (15) | 0.0000* | 0.0000 | 0.0000 | 0.3366 |
| **GAI** | 117,9 (12.2) | 65.6 (18.4) | 76.7 (14.6) | 0.0000* | 0.0000 | 0.0002 | 0.4457 |
| **WMI** | 110.5 (10.1) | 68 (11.6) | 76 (14.0) | 0.0000* | 0.0000 | 0.0002 | 0.5163 |
| **PSI** | 106.6 (16.3) | 62.6 (8,2) | 68.7 (13) | 0.0000* | 0.0001 | 0.0003 | 0.7471 |
| **CCAS** | | | | | | | |
| **Category switching** | 15 (12-15) | 9 (4-12) | 10 (8-15) | 0.0052§ | 0.0038 | 0.0215 | 0.2133 |
| **Go_nogo** | 1.5 (0-2) | 1 (0-1) | 0.5 (0-2) | 0.2083§ | | | |
| **Digit span backwards** | 6 (4-6) | 3 (2-5) | 2 (1-4) | 0.0016§ | 0.0257 | 0.0008 | 0.1582 |
| **Affectivity** | 6 (5-6) | 5 (2-6) | 4.5 (1-6) | 0.0694§ | | | |
| **ROCF** | 33.8 (21.8-35.7) | 27.7 (21.8-32.4) | 23.7 (15.2-30.5) | 0.0161§ | 0.0509 | 0.0098 | 0,2782 |
| **ToL** | | | | | | | |
| **Total Initiation Time** | 109.3 (17.5) | 104.4 (9.8) | 97.7 (8.4) | 0.3072* | | | |
| **Total Execution Time** | 99.8 (8.5) | 51.6 (33.5) | 71.4 (16) | 0.0018* | 0.0016 | 0.0436 | 0.2499 |
| **Total Correct score** | 106.5 (15.6) | 97.6 (9.21) | 93.7 (8.9) | 0.1677* | | | |

| | | | | | | | |
|---|---|---|---|---|---|---|---|
| **Total move score** | 103.5 (13.2) | 87.2 (16.6) | 88.4 (20.5) | 0.1599* | | | |
| **Time violations** | 107 (81-108) | 64 (10-106) | 80 (58-108) | 0.0146§ | 0.0081 | 0.0575 | 0.1775 |
| **Rule Violations** | 105 (74-106) | 78 (0-104) | 105 (55-106) | 0.1412§ | | | |
| **CPT-3** | | | | | | | |
| **Hit Reaction Time** | 43.9 (2.1) | 53.7 (8.34) | 53.8 (11.4) | 0.0455* | 0.0977 | 0.0734 | 0.9995 |
| **Attention Reaction Time** | 39.3 (5.1) | 61.7 (13.2) | 57.5 (8.5) | 0.0006* | 0.0011 | 0.0040 | 0.7185 |
| **Omissions** | 43 (43-48) | 59 (50-275) | 56 (45-90) | 0.0052§ | 0.0034 | 0.0244 | 0.1936 |
| **Commission errors** | 46 (36-62) | 75 (57-84) | 72 (49-78) | 0.0044§ | 0.0038 | 0.0151 | 0.2495 |
| **Perseveration** | 45 (45-48) | 47 (45-54) | 53 (48-90) | 0.0111§ | 0.3316 | 0.0058 | 0.0302 |
| **Vigilance** | 46 (43-49) | 30 (18-66) | 55 (33-64) | 0.1928§ | | | |
| **Sustained attention** | 47.9 (3.9) | 47.3 (11.5) | 55.7 (10.1) | 0.1861* | | | |
| **ABCL** | | | | | | | |
| **Anxious / Depressed** | 53.6 (2.5) | 59.6 (6.9) | 59.7 (6.4) | 0.0737* | | | |
| **Withdrawn** | 51 (50-63) | 54 (51-68) | 68 (50-81) | 0.0662§ | | | |
| **Somatic Complaints** | 57.5 (6.7) | 56.2 (6.8) | 54.5 (5.4) | 0.6894* | | | |
| **Internalizing Problems** | 53.4 (6.3) | 57.8 (8.3) | 60.2 (7.8) | 0.2416* | | | |
| **SRS-2- SCI** | 42 (8.4) | 56.8 (9.1) | 54.8 (14.3) | 0.0440* | 0.0685 | 0.0998 | 0.9511 |

SARA (Scale for the Assessment and Rating of Ataxia, administered only in JS and SP); CCAS (Cerebellar cognitive affective syndrome scale); WAIS (Wechsler adult intelligence scale); FSIQ (Full Scale Intelligence Quotient), GAI (General Ability Index); WMI (Working Memory Index); Processing Speed Index (PSI); ROCF (Rey–Osterrieth Complex Figure); Tol (tower of London); CPT-3 (continuous performance test 3; ABCL (Adult behaviour checklist), and SRS_SCI ) Social Responsiveness Scale - Social Communication and Interaction index).

**Supplementary Figure 1: Extended linear regression analyses.**

Scatter plots illustrating observed versus predicted values for all additional stepwise linear regression models reported in Fig. 7 of the main text. Each plot represents one regression model linking MRI-derived anatomo-physiological parameters to a specific clinical or neuropsychological score, stratified by functional network. Each point corresponds to a single subject, with colours indicating group affiliation. The grey dashed line (y = x) represents the line of perfect prediction, while the red line indicates the regression fit of the predicted values. These plots complement the examples shown in the main text by providing a complete visualisation of model performance across all significant associations.

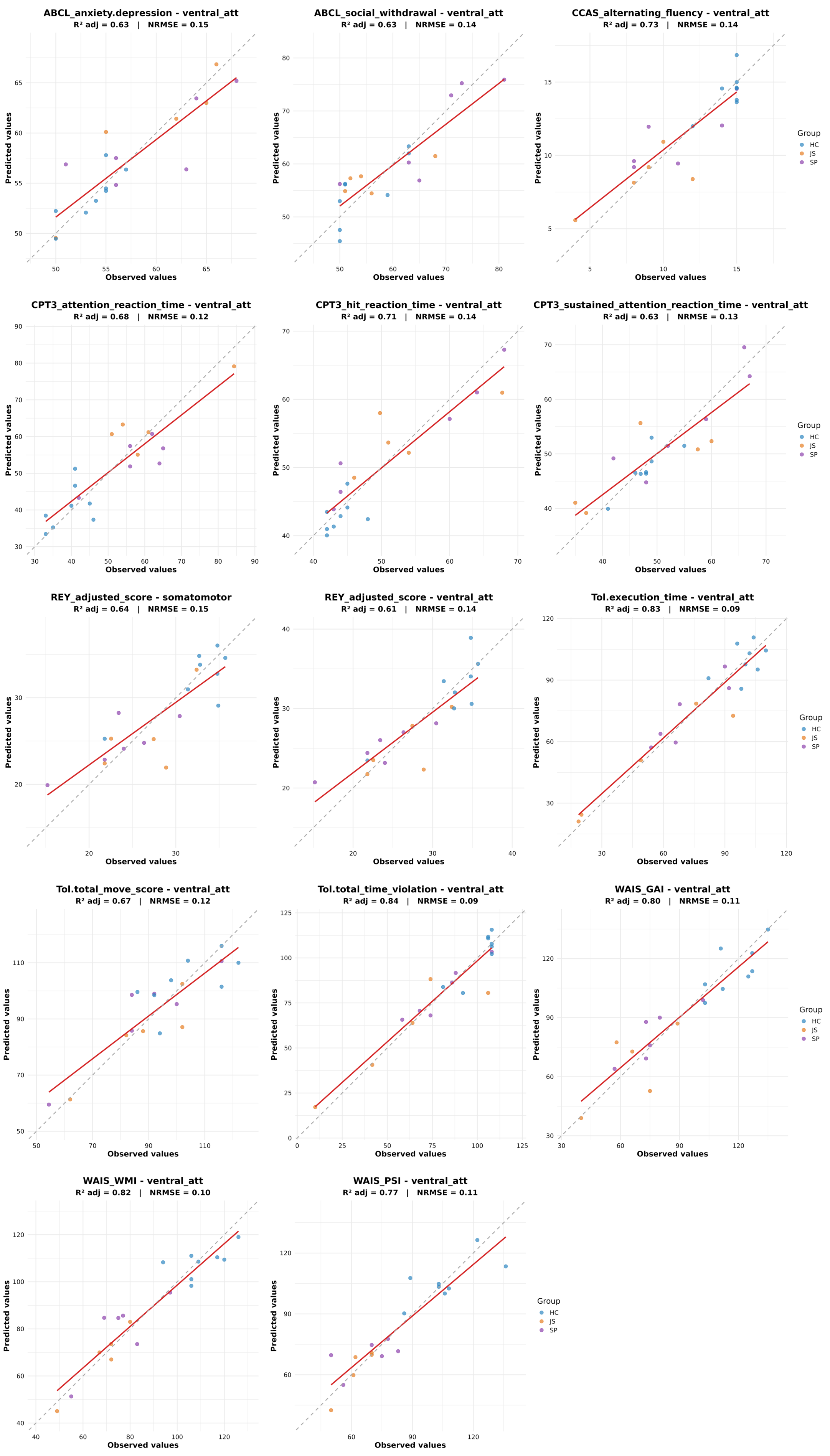

ABCL_anxiety.depression - ventral_att
R² adj = 0.63 | NRMSE = 0.15
ABCL_social_withdrawal - ventral_att
R² adj = 0.63 | NRMSE = 0.14
CCAS_alternating_fluency - ventral_att
R² adj = 0.73 | NRMSE = 0.14
CPT3_attention_reaction_time - ventral_att
R² adj = 0.68 | NRMSE = 0.12
CPT3_hit_reaction_time - ventral_att
R² adj = 0.71 | NRMSE = 0.14
CPT3_sustained_attention_reaction_time - ventral_att
R² adj = 0.63 | NRMSE = 0.13
REY_adjusted_score - somatomotor
R² adj = 0.64 | NRMSE = 0.15
REY_adjusted_score - ventral_att
R² adj = 0.61 | NRMSE = 0.14
Tol.execution_time - ventral_att
R² adj = 0.83 | NRMSE = 0.09
Tol.total_move_score - ventral_att
R² adj = 0.67 | NRMSE = 0.12
Tol.total_time_violation - ventral_att
R² adj = 0.84 | NRMSE = 0.09
WAIS_GAI - ventral_att
R² adj = 0.80 | NRMSE = 0.11
WAIS_WMI - ventral_att
R² adj = 0.82 | NRMSE = 0.10
WAIS_PSI - ventral_att
R² adj = 0.77 | NRMSE = 0.11
Predicted values
Observed values
Group
HC
JS
SP
